# Encapsulation boosts islet-cell signature in differentiating human induced pluripotent stem cells via integrin signalling

**DOI:** 10.1101/791442

**Authors:** Heidrun Vethe, Thomas Aga Legøy, Shadab Abadpour, Berit L. Strand, Hanne Scholz, Joao A. Paulo, Helge Ræder, Luiza Ghila, Simona Chera

## Abstract

Cell replacement therapies hold great therapeutic potential. Nevertheless, our knowledge of the mechanisms governing the developmental processes is limited, impeding the quality of differentiation protocols. Generating insulin-expressing cells *in vitro* is no exception, with the guided series of differentiation events producing heterogeneous cell populations that display mixed pancreatic islet phenotypes and immaturity. The achievement of terminal differentiation ultimately requires the *in vivo* transplantation of, usually, encapsulated cells. Here we show the impact of cell confinement on the pancreatic islet signature during the guided differentiation of alginate encapsulated human induced pluripotent stem cells (hiPSCs). Our results show that encapsulation improves differentiation by significantly reshaping the proteome landscape of the cells towards an islet-like signature. Pathway analysis is suggestive of integrins transducing the encapsulation effect into intracellular signalling cascades promoting differentiation. These analyses provide a molecular framework for understanding the confinement effects on hiPSCs differentiation while confirming its importance for this process.

## Introduction

Differentiation of human pluripotent stem cells (hPSCs) into fully developed cell types holds a great therapeutic potential, especially the generation of hPSC-derived insulin-producing cells for cell therapy of diabetes. In mammals, pancreatic adult islets contain four hormone-secreting endocrine cell populations, each secreting only one specific hormone: insulin (β-cells), glucagon (α-cells), somatostatin (δ-cells), pancreatic polypeptide (PP-cells, γ-cells) and ghrelin (ɛ-cells). Multistep-protocols that mimic pancreas development have been designed for the generation of hPSC-derived insulin-expressing cells *in vitro*^1–3^.

Despite an increasing number of differentiation protocols, the efficient generation of a stable mature and functional insulin-producing β-cell is yet to be convincingly achieved4,5. Moreover, most of these protocols do not restrict the cells towards β-cell fate exclusively, but rather generate cell entities resembling diverse hormone-secreting islet cell types. At final stages of differentiation, the generated cell populations are highly heterogeneous, with cells displaying variable levels or absence of key β-cell markers (such as PDX1 and NKX6.1) and a polyhormonal profile, resembling the transient endocrine cells observed in the fetal pancreas^6,7^. Nevertheless, the reported heterogeneity grants certain advantages, as increasing evidence indicates that efficient glucose regulation requires proper arrangement and coordination between the various endocrine cell types found within the islets^8,9^. Therefore, recent research suggested focusing on *in vitro* generation of islets (*i.e.* all five cell populations) rather than aiming at differentiating cells into a single type of specialized cells^10^.

Currently, the final steps of β-cell maturation are achieved *in vivo*, by transplantation into mammalian hosts, such as mice. The cellular and molecular basis of the process driving this *in vivo* terminal differentiation is not yet completely understood^11^. Different potential scenarios include the involvement of circulating factors^12–15^, nervous system association^16–18^ and the presence of a 3D niche^19,20^, amongst others. Discriminating the exact contribution of each of these potential scenarios on the transplanted hPSC-derived cells is difficult due to the inherent complexity of the organism environment.

Microencapsulation of islets into alginate microbeads was used first in the 1980s^21^, and was later employed in several studies for transplantation of pancreatic islets^22–25^. Previous studies have reported that entrapment of hPSCs under the 3D environment of alginate microcapsules^26^ supports long-term maintenance of pluripotency^27^ and differentiation of dopamine neurons^28^, as well as pancreatic progenitors^29^. Alginate is recognized for properties and characteristics such as its ability to make hydrogels at physiological conditions, transparency for microscopic evaluation, gel pore network that allows diffusion of nutrients and waste materials^30^, making alginate an attractive alternative for embedding hPSC-derived cells during differentiation.

In this study, we differentiated hiPSCs (human induced pluripotent stem cells) towards β-like cells following a seven-stage protocol^1^, as we have reported previously^31^, to assess the impact of alginate encapsulation on islet cell differentiation potential during *in vitro* differentiation. Our data indicate that encapsulation of pancreatic endocrine progenitor efficiently improves the differentiation outcome by increasing both the proportion of hormone-positive cells and the fraction of insulin cells co-expressing key β-cell markers. Moreover, encapsulation enables proteome adaptations of the differentiating cells towards a more islet-like fingerprint in a stage-specific manner, where the encapsulation of the first differentiation stages promotes early differentiation signals, while the encapsulation at a later differentiation stage promotes hormones and factors involved in hormone synthesis and secretion. Our results further suggest that these effects of alginate are relayed through integrins, which presumably translate the pressure elicited by the confinement of cells in the alginate matrix into signalling cascades.

## RESULTS

### Encapsulation promotes the expression of islet hormones and key islet transcriptional regulators

To investigate whether encapsulation had an impact on the differentiation outcome, we differentiated cells either on Matrigel-coated plates (representing a classical 2D culture condition) or encapsulated in alginate (representing a 3D platform for differentiation). Due to its high reproducibility and feasibility, we selected one of the most commonly employed protocol for β-cell differentiation designed by Rezania et al^1^, where hiPSC cells are differentiated gradually through a sequence of events involving 7-stages (Supp. Figure 1a).

The dissociated cells before encapsulation displayed a viability of 90.96%±5.6, presenting homogeneous cell morphologies and no signs of proliferation or cluster formation within the alginate beads. Sporadic clumps of cells escaping the dissociating procedure were observed, however these did not present an ulterior different behaviour compared to single cells (Supp. Figure 1b, c). The alginate beads, presenting a radius of 375-500µm, were processed for immunofluorescence (IF) and high magnifications of whole mount encapsulated cells (Supp. Figure 1c) and sections (Supp. Figure 1d) revealed spherical cell morphology typical of single cells in suspension. To allow the quantification of the largest possible bead volume, we imaged for each whole mount bead a mosaic of 3×3 fields of view (FOV) over 100 different focal planes and performed a 3D reconstruction (Supp. Figure 1e, f). Due to the large amount of data generated and also for excluding any potential counting-bias, we performed automatic quantification using Imaris 9.1.2 with the program being supervised by manual counting of randomly chosen FOV.

Cells were encapsulated: (1) at the beginning of the protocol (hiPSC stage [S0]) and differentiated in alginate capsules until Stage 7 (hereafter termed S7^bead[S0-S7]^), (2) during the last two stages of the differentiation protocol, i.e. end of Stage 5 (corresponding to pancreatic endocrine progenitors) until Stage 7 (hereafter termed S7^bead[S5-S7]^) as well as (3) at the final stage of the differentiation protocol, just before fixation, for consistency of imaging and counting (hereafter termed S7^bead[S7]^) (Figure 1a, Supp. Figure 1a, b).

**Figure 1.**
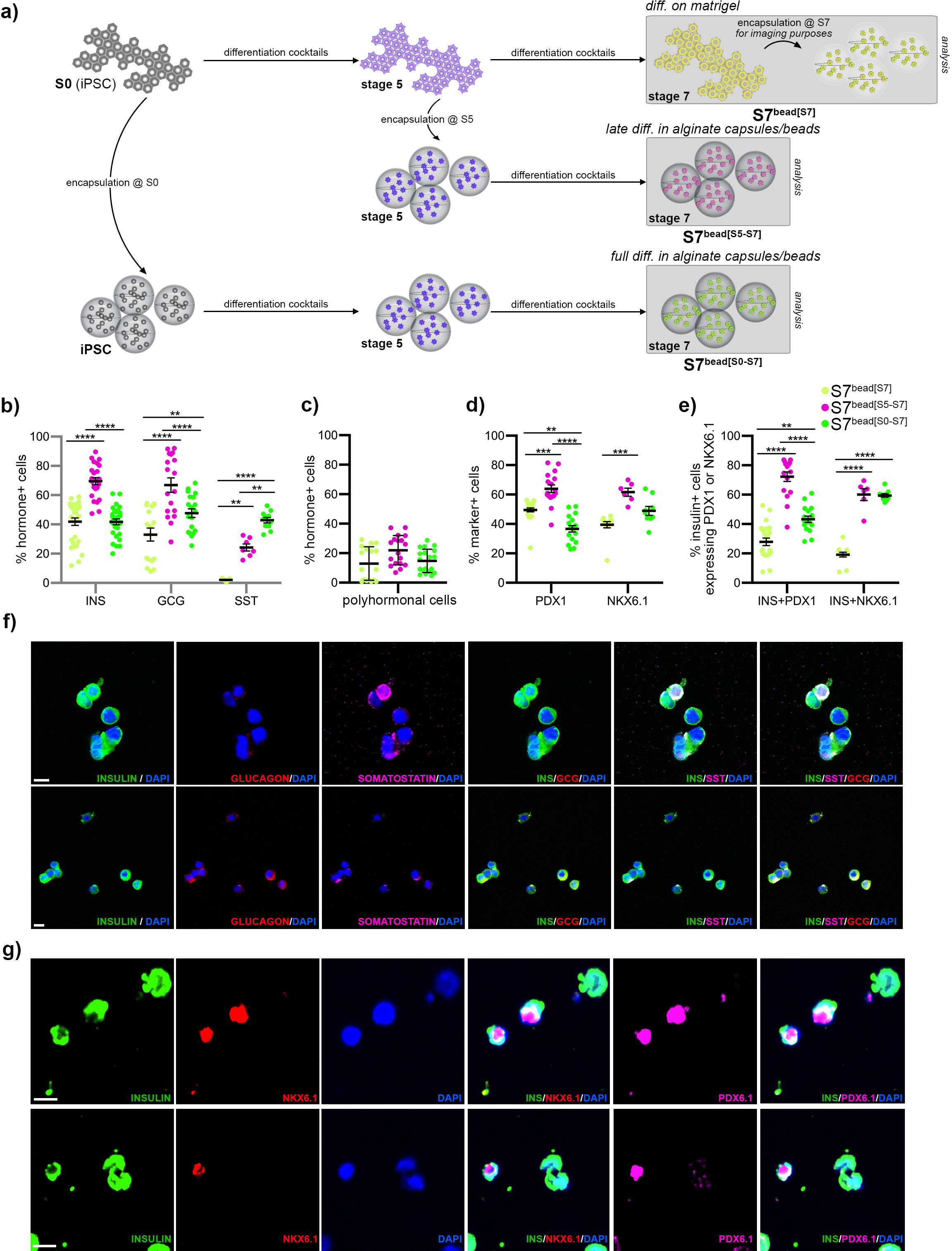
Comparison of the hiPSC differentiation outcome according to the stage of encapsulation. a) Scheme depicting the three cell populations analysed by immunofluorescence. b) Proportion of the differentiated hiPSC-cells expressing insulin, glucagon or somatostatin in the three distinct populations analysed, quantified by Imaris software. c) Proportion of bihormonal cells in the three distinct populations analyzed. d) Proportion of the differentiated hiPSC-cells expressing PDX1 or NKX6.1 in the three distinct populations analysed. e) Proportion of insulin+ cells coexpressing PDX1 or NKX6.1. f) High magnification confocal images of cells inside alginate capsules stained for insulin (green), glucagon (red), somatostatin (purple) and DAPI (blue) by whole mount immunofluorescence. g) Whole mount immunofluorescence of encapsulated cells stained for insulin (green), NKX6.1 (red), PDX1 (purple) and DAPI (blue), gamma correction 0.4. Scale bars: 10µm. Graphs data are shown as mean ± SEM.

We first assessed the variation in the proportion of insulin-, glucagon-, and somatostatin-expressing cells following encapsulation (Figure 1b, c, f, Supp. Figure 2a). A highly significant increase in the proportion of cells expressing insulin, glucagon, and somatostatin was observed when cells were encapsulated at the later stages of differentiation (S7^bead[S5-S7]^) as compared to the ones grown on Matrigel-coated plates (S7^bead[S7]^). The number of bihormonal cells was not significantly affected by encapsulation (Figure 1c). Interestingly, the cells encapsulated during the entire period of differentiation (S7^bead[S0-S7]^) displayed a significant increase of the glucagon- and somatostatin-expressing cell populations, but not insulin, as previously reported^29^.

We further assessed in all three conditions analysed the expression of the β-cells transcriptional regulators PDX1 and NKX6.1 (Figure 1d, e, g, Supp. Figure 2b), two key factors involved in early pancreas development, endocrine compartment differentiation and β-cell fate^32–34^. Consistent with the previous results, the highest increase in the proportion of the PDX1- and NKX6.1-expressing cells was observed in the S7^bead[S5-S7]^ population (63.97%, Figure 1c, e). The proportion of PDX1-expressing cell population was significantly decreased in the S7^bead[S0-S7]^ (36.7%) when compared to the S7^bead[S7]^ population differentiated in 2D (49.53%), while the proportion of NKX6.1-expressing cells was not significantly different. Despite the higher proportion of PDX1-expressing cells, only a very low number of the insulin-positive cells of S7^bead[S7]^ population co-expressed PDX1 (27.89%) and even fewer (19.19%) co-expressed NKX6.1 (Figure 1e, g, Supp. Figure 2b) indicating the presence of (1) a large fraction of insulin-expressing cells missing these key factors for their functionality and stability as well as (2) a considerable, probably immature, insulin-negative subpopulation of PDX1+ and NKX6.1+ cells.

In contrast, despite the lower proportion of PDX1-expressing cells, the S7^bead[S0-S7]^ had a higher proportion of insulin-positive cells co-expressing PDX1 (43.33%) as well as NKX6.1 (59.44%). The best expression overlap was identified once more in the population of cells encapsulated during the last two stages of differentiation (S7^bead[S5-S7]^) with 72.25% of the insulin+ cells co-expressing PDX1 and 60.04% co-expressing NKX6.1 (Figure 1e, g, Supp. Figure 2b).

Overall, these data indicate that encapsulation during the last stages of differentiation (*i.e.* from the pancreatic endocrine progenitor stage [S5]) improves the differentiation outcome by increasing both the proportion of hormone-positive cells and the fraction of insulin cells co-expressing key β-cell markers. Interestingly, encapsulation during the entire length of differentiation seem to improve the insulin cells profile quality rather than increasing their numbers.

### Encapsulation promotes a large battery of proteins towards islet-like abundance levels

To comprehensively assess the encapsulation effect on the differentiation outcome, in a parallel and independent set of experiments, we performed global proteomics on differentiating hiPSC cells, which were grown either on Matrigel-coated plates (2D) or encapsulated in alginate (3D). To allow the investigation of the observed stage-dependent encapsulation effect, the cells were differentiated in capsules during (1) the early stages of differentiation (from the iPSC stage [S0] to the pancreatic endocrine progenitor [S5], hereafter termed S5^bead[S0-S5]^) and (2) the late stages of differentiation (from the pancreatic progenitor stage [S5] to maturing β-like cell [S7], hereafter termed S7^bead[S5-S7]^) (Figure 2a). These, their Matrigel-differentiating counterparts (S5 and S7) and native human islets isolated from deceased donors were compared by TMT 11-plex-based quantitative proteomics (Figure 2b). We quantified a total of 5364 proteins and focused our analysis on the proteins expressed in at least one sample of each condition (5029 proteins).

**Figure 2.**
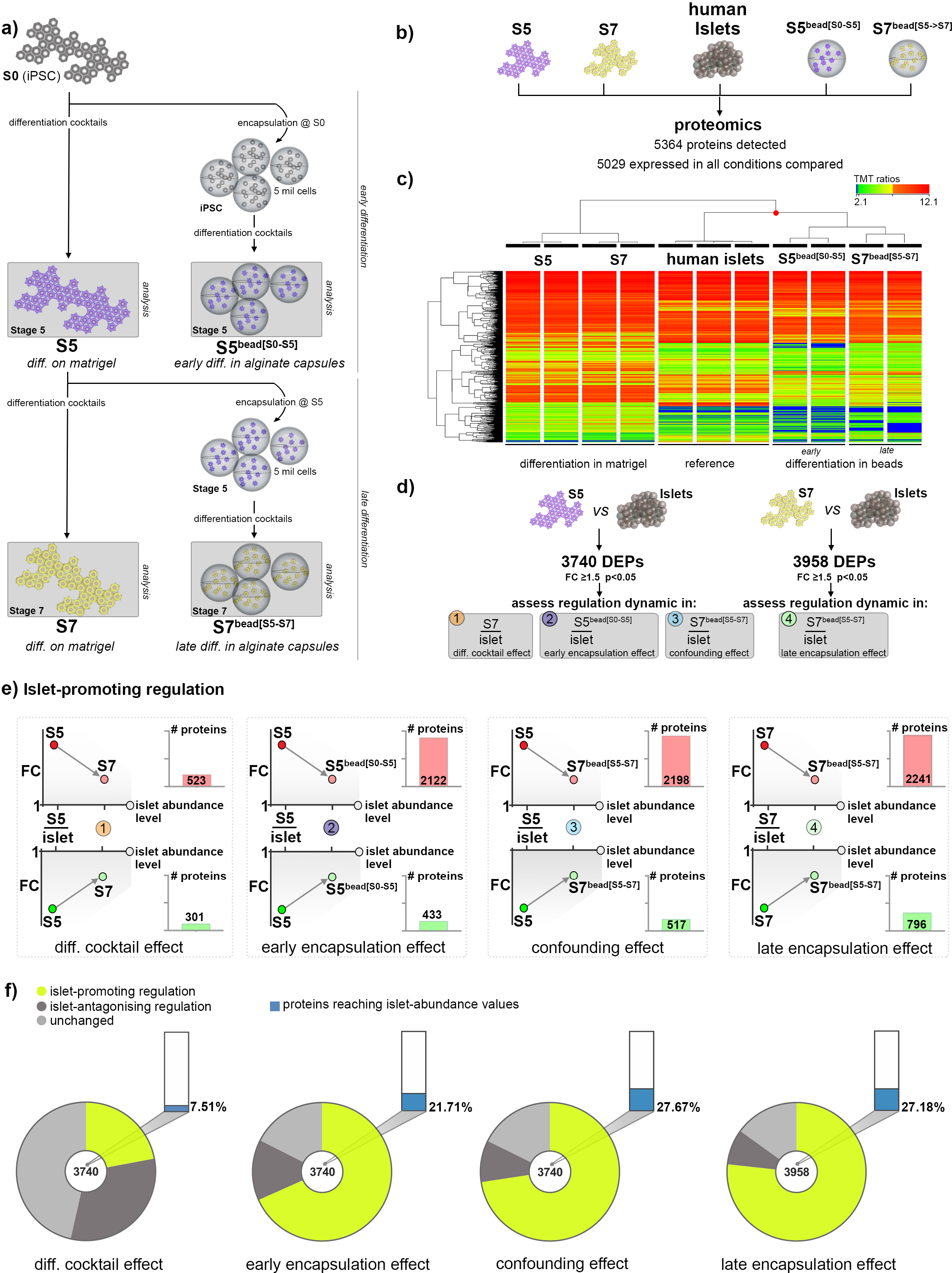
Global proteome analysis of hiPSC differentiating either on Matrigel or encapsulated in alginate capsules. a) Scheme illustrating the cell populations and differentiation stages considered for global proteomics. b) Experimental design of the conditions compared in TMT 11-plex proteomics. c) Hierarchical clustering of normalized TMT-ratios (n=2,2,2,2,3). d) Analysis workflow depicting the comparisons employed and the assessed corresponding effect. e) The number of proteins showing a dynamic of regulations compatible with an islet-promoting pattern in response to each of the four effects considered. Arrows depict the generic prerequisite direction of regulation for group inclusion. f) Pies charts depicting the proportion of proteins following islet-promoting and islet-antagonizing regulation patterns in each of the four effects considered. The graph bars represent the proportion of proteins reaching abundance levels indistinguishable from those detected in native human islets.

The hierarchical clustering revealed that encapsulated samples (S5^bead[S0-S5]^ and S7^bead[S5-S7]^) cluster closer to the human islets than their 2D culture condition differentiation-stage counterparts, with the S7^bead[S5-S7]^ samples being nearest (Figure 2c). These data suggest that encapsulation globally enhances the pancreatic endocrine cell fate, reinforcing the initial IF results.

Unfortunately, due to the inherent sensitivity limitation of the proteomics methods, we were unable to detect proteins with very low abundance, as displayed by many key pancreatic islet transcription factors, such as PDX1 or NKX6.1. Nevertheless, in consensus with our previous IF characterization, we observed a steep increase of all pancreatic islet hormones following encapsulation (Supp. Figure 3a).

We further aimed to identify the proteins modulated towards an islet-like regulation in response to encapsulation. A simple comparison analysis between cells differentiating in capsules and their 2D culture counterparts would provide information about the general effects of encapsulation, but would fall short in identifying proteins relevant for differentiation towards islet cell phenotypes. Consequently, to discriminate the differentially-expressed proteins (DEPs) that in response to encapsulation are changing their abundance profiles towards the levels detected in islets from those with opposite regulation, we introduced the islet samples as normalization. We first filtered 3740 DEPs (FC≥1.5 p<0.05) between S5-cells and islets (Supp. Table 1) and then assessed their regulation dynamics in response to the different conditions (Figure 2d, outlined in Supp. Figure 3b): (1) stages of the differentiation protocol (S7/Islet, hereafter termed “Differentiation Cocktail Effect”), (2) encapsulation during early stages of differentiation (S5^bead[S0-S5]^/Islet, hereafter termed “Early Encapsulation Effect”) and (3) differentiation cocktail combined with late encapsulation (S7^bead[S5-S7]^/Islet, hereafter “Confounding Effect”). Following the same rationale, a second 3958 DEPs list (FC≥1.5 p<0.05) was generated between S7-cells and islets (S7/Islets, Supp. Table 1) and their regulatory dynamics in response to (4) encapsulation during later stages of differentiation was assessed (S7^bead[S5-S7]^/Islet, hereafter termed “Late Encapsulation Effect”) (Figure 2d, Supp. Figure 3b).

We categorized the DEPs according to their response to each of the above effects (up- or downregulated, FC≥1.5 compared to their initial level of regulation in S5/Islet or S7/Islet, respectively, Figure 2e, Supp. Figure 3c). The upregulated DEPs (S5/Islet or S7/Islet), which decrease their abundance towards islet expression levels following a certain effect (differentiation cocktail, early encapsulation, confounding) present an islet-promoting regulation (Figure 2e, upper row). Following the same rationale, downregulated DEPs that show a tendency to recover towards islet expression levels will fall into the same category (Figure 2e, lower row). In contrast, proteins that respond by drifting away from the islet abundance levels, will display an islet-antagonizing regulation (Supp. Figure 3c). The number of proteins with an islet-promoting regulation was steeply increased in response to encapsulation when compared with the effect of the differentiation cocktail, regardless of the time of alginate encapsulation (Figure 2e, f). Most of these proteins (83%, 81%, 73.8%, depending on the effect) presented a pattern of regulation characterized by high abundance in S5- or S7-cells as compared with islets, which decreased towards islet expression levels upon encapsulation (Figure 2e, compare red and green bars of the last three effects).

Furthermore, the number of proteins reaching abundance values similar to those exhibited by native islets was also increased by encapsulation, from 7.51% regulated in response to the differentiation cocktail alone to 21.71%, 27.67% and 27.18% when encapsulation influence was considered (Figure 2f, Supp. Table 2). Of note, the number of proteins presenting an islet-antagonizing pattern was decreasing in the three effects assessing the encapsulation involvement (Figure 2f dark grey sectors, Supp. Figure 3c).

Overall, these results suggest that, regardless of the iPSC differentiation stage, encapsulation was able to substantially modify the proteome landscape towards a more islet-like fingerprint, mostly by either maintaining or promoting lower abundance levels of certain proteins, which otherwise display an upregulated pattern in 2D-differentiated cells.

### Pathway analysis reveals common proteome signatures in response to encapsulation

To identify the protein networks, signalling pathways and upstream regulators characterizing the improved islet-signature of the encapsulated differentiating cells in each of the four effects described above, we performed pathway analysis on the protein sets exhibiting islet-promoting regulation (Figure 3, Supp. Figure 4a, c, e, g, Supp. Table 2). As a further refinement, we repeated the analysis on the protein sets reaching islet-like abundance levels in each dataset (Supp. Figure 4b, d, f, h).

**Figure 3.**
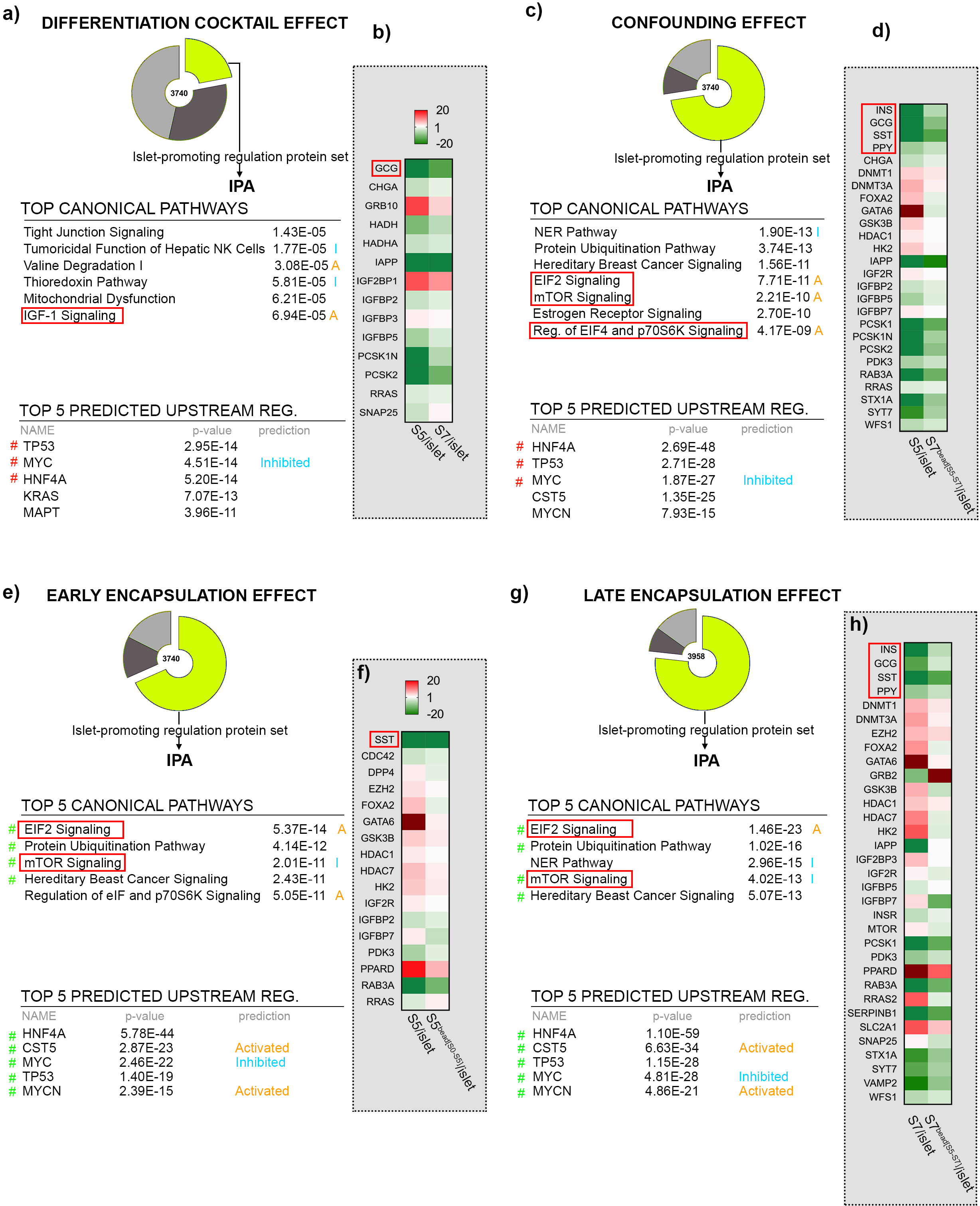
Pathway analysis and heatmaps of proteins following islet-promoting regulation patterns in response to the different effects assessed. a) Tables depicting the top canonical pathways and predicted upstream regulators in response to Differentiation Cocktail Effect, b) Heatmap representing the regulation of selected markers in response to Differentiation Cocktail Effect c) Tables depicting the top canonical pathways and predicted upstream regulators in response to Confounding Effect, d) Heatmap representing the regulation of selected markers in response to Confounding Effect, e) Tables depicting the top canonical pathways and predicted upstream regulators in response to Early Encapsulation Effect, f) Heatmap representing the regulation of selected markers in response to Early Encapsulation Effect, g) Tables depicting the top canonical pathways and predicted upstream regulators in response to Late Encapsulation Effect, h) Heatmap representing the regulation of selected markers in response to Late Encapsulation Effect (orange A – predicted activation, blue I – predicted inhibition, red # signals shared pathways or predicted upstream regulators between left and right effects).

IPA (Ingenuity Pathway Analysis software) revealed that the protein set exhibiting an islet-promoting pattern of regulation in response to the differentiation cocktail (Differentiation Cocktail Effect) was consistent with the expected outcome of the typical stage progression of differentiating cells (Figure 3a, b, Supplemental Figure 4a). The Tight Junction Signaling, several metabolic pathways and IGF-1 Signaling were identified in the top canonical pathways, the latter being predicted as activated in S7-cells (Figure 3a). The program selected the cell-cycle regulators TP53, MYC, KRAS as well as the key developmental pancreatic islet cell marker HNF4A and the insulin trafficking regulator MAPT (TAU protein) in the Top 5 predicted upstream regulators responsible for the observed proteome landscape. Moreover, MYC was predicted as inhibited, in agreement with the expected low proliferation rate exhibited by differentiated 7-cells (Supp. Figure 4a). Of note, three of the top upstream regulators (TP53, MYC and HNF4A) were predicted in the Top5 of all four effects, with the prediction of MYC inhibition being also a shared feature.

In addition, the protein set contained key markers of pancreatic-islet cells, such as the hormones GCG (glucagon) and IAPP (Islet Amyloid Polypeptide, co-secreted with insulin by the β-cells), the hormone processing proconvertases PCSK1N and PCSK2 (alpha-cell marker) as well as the pan-endocrine marker CHGA (chromogranin A) (Figure 3b).

The pathway analysis of the protein subset attaining abundance values similar to islets identified IGF-1 Signaling as being the Top 3 canonical pathway and HNF4A as top predicted upstream regulator, suggesting that at least some of the markers governing islet cell differentiation reached the desired islet-like abundance (Supplemental Figure 4b). Taken together, these results indicate an expected improved differentiation status of the cells following the addition of the differentiation cocktails. Of note, the action of the differentiation protocol cocktails seems to mostly promote α-cell fate markers, a problem already described by previous studies^35^.

For the Confounding Effect (*i.e.* the observed modulation of the proteins could not be unequivocally attributed to the differentiation cocktail, late encapsulation or both) the pathway analysis of the protein set exhibiting an islet-promoting pattern of regulation, revealed the involvement of the PI3K/AKT axis and mTOR signalling within the Top 5 canonical pathways (Figure 3c). The proteome regulatory signature was consistent with the activation of important upstream regulators for pancreatic-islet differentiation such as SOX4 and MAF (Supp. Figure 4c). Moreover, the Confounding effect prompted the islet-promoting regulation of all standard islet hormones, including insulin, as well as key molecular components involved in β-cell function and generally in hormone synthesis and secretion (Figure 3d). The analysis of the proteins reaching islet-abundance values identified similar canonical pathways, with seven of the top pathways involving components of PI3/AKT or mTOR signalling (Supp. Figure 4d).

Interestingly, the Top 5 canonical pathways were overlapping with the Late Encapsulation Effect and partially overlapping with the Early Encapsulation Effect, with slight differences in rank (Figure 3c, e, g). Moreover, the Early and Late Encapsulation effects share their prediction of Top 5 upstream regulators. Last, the Late Encapsulation Effect presents a similar regulation of all islet hormones (Figure 3h). Overall, these data suggest that the encapsulation and not the differentiation cocktail was the driving-force promoting the improved islet-signature regulation of the Confounding effect.

The pathway analysis of the encapsulation-based effects (Early and Late Encapsulation effects), showed a high degree of overlap for both the Top 5 canonical pathways and the Top 5 predicted upstream regulators (4/5), displaying the involvement of the PI3K/AKT axis and mTOR Signalling as well as HNF4A and CST5 as top upstream regulators (Figure 3e, g). The proteome signatures of both effects allowed the prediction of CDKN2A (p16) activation as well as MYC1 and FOXM1 inhibition, suggesting a decrease in the differentiating cell proliferation in response to encapsulation, regardless of the encapsulation stage (Supp. Figure 4e, g). Specific for the Early Encapsulation Effect, the proteome profile allowed the prediction of PDX1 activation as well as the inhibition of β-catenin (Supp. Figure 4e), suggesting a differentiation event promoted by encapsulation during the early stages of differentiation. The Early Encapsulation Effect promoted the regulation of only one islet hormone, SST (somatostatin), as well as chromatin modifiers (such as EZH2) and β-cell differentiation factors, consistent with the early differentiation stage of these cells (Figure 3f).

In contrast, the Late Encapsulation Effect induced the islet-promoting regulation of all pancreatic islet hormones (Figure 3h). Besides hormones, a large number of key factors involved in hormone synthesis and secretion were regulated towards islet-abundance values, including the main pancreatic islet-cells proconvertases PCSK1 (β-cell marker) and the main glucose transporter in humans, GLUT1 (SLC2A1) (Figure 3h). As expected, the analysis of the protein subset displaying islet-like abundance values reinforced the involvement of the PI3K/AKT axis and mTOR Signaling in addition to HNF4A as a top predicted upstream regulator (Supp. Figure 4f, h).

In conclusion, the encapsulation during the late stages of differentiation promotes the pancreatic-islet differentiated profile in S7-cells, exhibiting increased level of hormones and hormone proconvertases.

Overall, these results suggest that the Differentiation Cocktail Effect presented the most divergent signature of the protein set with islet-promoting regulation, while the three effects involving cell encapsulation exhibit rather related signatures. Moreover, the effect of encapsulation seems to have a stronger impact on the proteome profile of cells than the effect of the differentiation cocktail, as inferred from the Confounding Effect analysis. Last, the Early Encapsulation Effect seems to be able to promote early differentiation signals towards islet cell fate, while the Late Encapsulation effect promotes hormones and factors involved hormone synthesis and secretion, hence boosting the differentiated islet-cell profile.

### Encapsulation and the differentiation cocktail promote different protein sets

To identify whether encapsulation acts on the same targets as the differentiation cocktail, we first filtered for proteins exhibiting islet-promoting regulation shared by Differentiation Cocktail Effect, Early Encapsulation Effect and Late Encapsulation Effect (Figure 4a, left panel, Supp. Table 3). We identified only 210 proteins as being regulated by all three effects, representing 25.5%, 8.2% and 6.9% of the respective protein sets, with the pathway analysis retrieving mainly metabolic pathways.

**Figure 4.**
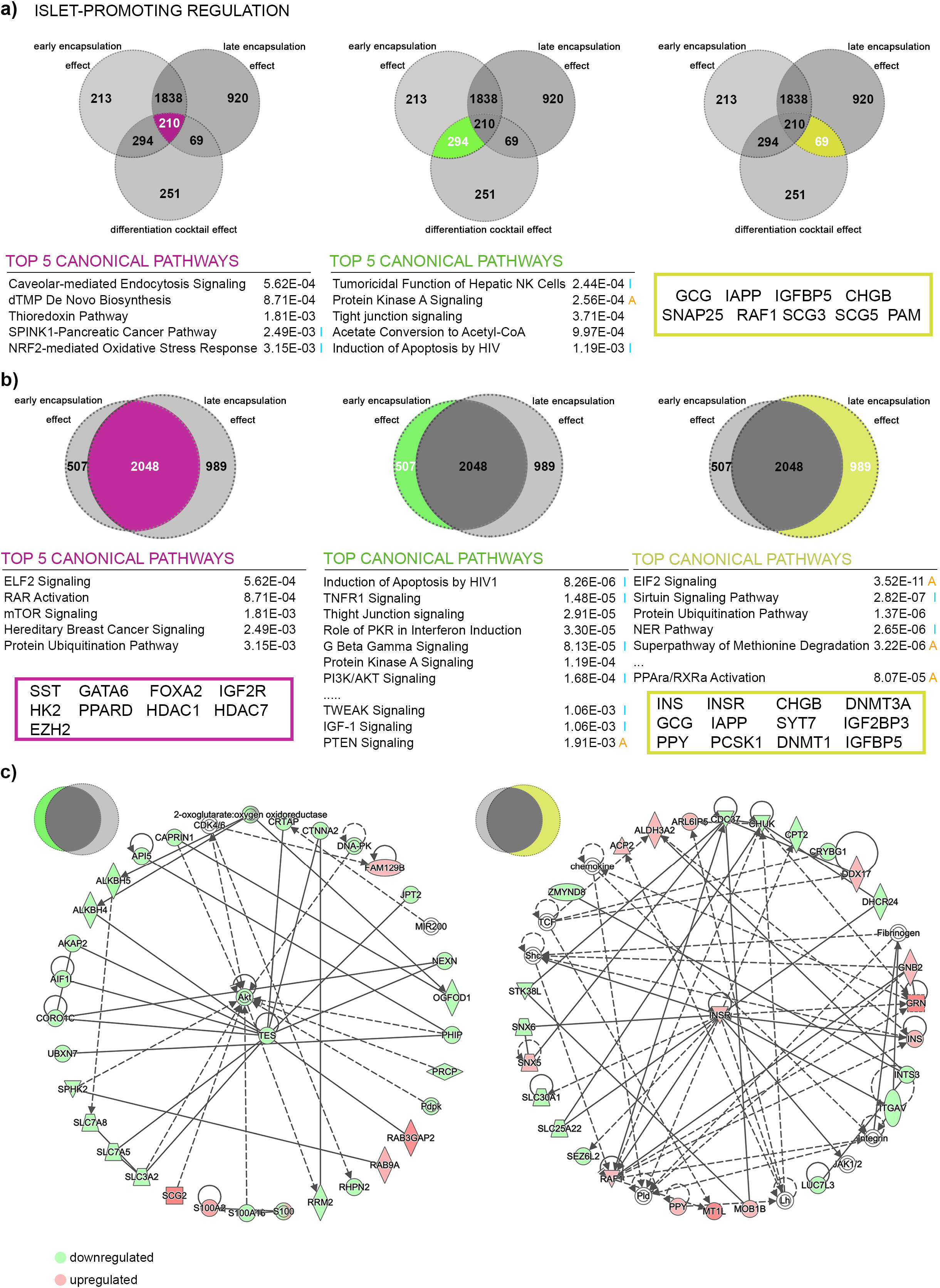
Pathways analysis of proteins displaying islet-promoting regulation patterns in response to more than one effect. a) IPA-generated tables of the Top 5 canonical pathways for proteome landscapes regulated in response to all three effects considered (purple, left Venn diagram) as well as to Differentiation Cocktail Effect and Early Encapsulation Effect solely (green, middle Venn diagram). Selected regulated pancreatic islet markers are shown for the proteome landscape responding to Differentiation Cocktail Effect and Late Encapsulation Effect solely (yellow, right Venn diagram). b) IPA-generated tables of top canonical pathways and selected regulated pancreatic islet markers for proteome landscapes regulated by encapsulation regardless of differentiation stage of encapsulation (purple, left Venn diagram), only by encapsulation during the early stages (S0->S5) of differentiation (green, middle Venn diagram) and only by encapsulation during the late stages (S5->S7) of differentiation (yellow, right Venn diagram). c) Selected IPA circular networks for the proteome regulated exclusively by either Early Encapsulation Effect (left) or Late Encapsulation Effect (right). The heatmap represents the direction of regulation towards islet-abundance values in S5^bead[S0-S5]^ compared to S5 (left) and S7^bead[S5-S7]^ compared to S7 population. Orange A – predicted activation, blue I – predicted inhibition, green – observed downregulation, red – observed upregulation. The circular networks and the top canonical pathways were generated through the use of IPA (QIAGEN Inc., https://www.qiagenbio-informatics.com/products/ingenuity-pathway-analysis).

The same Venn comparison revealed 294 proteins being regulated exclusively by the Differentiation Cocktail and Early Encapsulation effects (Figure 4a, middle panel, Supp. Table 3). In this subset, IPA identified Protein Kinase A Signalling as well as Tight Junction Signalling in the Top 5 canonical pathways, with the first being predicted as activated. The third group, solely regulated by the Differentiation Cocktail and Late Encapsulation effects, contained 69 proteins (Figure 4a, right panel, Supp. Table 3). The pathway analysis strength was limited due to the very low number of proteins in this profile, however it contained several key pancreatic-islet markers among which GCG, IAPP and CHGB (chromogranin B).

These data indicated that the encapsulation and the differentiation cocktail largely regulate different targets. The encapsulation during early stages of differentiation shared the largest number of proteins exhibiting islet-promoting regulation with the effect of the differentiation cocktail, suggesting once more that encapsulation alone can, at least in some extent, promote the pancreatic islet-cell differentiation. Furthermore, the boost of key mature islet-cell markers induced by the differentiation cocktail reinforces the role of encapsulation during the later stages of differentiation in boosting the differentiated islet-cell signature.

### Encapsulation effects on differentiation are stage-specific

To compare further the Early and Late Encapsulation effects, we first performed pathway analysis on the shared group of proteins exhibiting islet-promoting regulation (Figure 4b, left panel, Supp. Table 3). The two effects share a large battery of proteins (2048) accounting for 80.16% and 67.43% of the respective protein sets. Pathway analysis revealed the EIF2 Pathway, RAR activation and mTOR Signaling as the Top 3 canonical pathways. Moreover, both effects collectively promoted towards islet-like levels several key pancreatic islet markers, including: the SST hormone, the FOXA2 and GATA6 developmental factors (central role in early pancreatic islet-cell differentiation) and the epigenetic modifiers EZH2 (histone methyltransferase, a key regulator of β-cell proliferation and regeneration), HDAC1 and HDAC7 (histone deacetylases with role in β-cell mass regulation). These results indicate that most of the proteins respond to encapsulation regardless of the encapsulation stage. Also, the presence of key pancreatic-islet epigenetic modifiers in this data set confirms once more the capacity of encapsulation to profoundly affect and promote islet-cell fate.

The pathway analysis of proteins exhibiting islet-promoting regulation triggered solely by early encapsulation (Figure 4b, middle panel, Supp. Table 3) indicated PI3K/AKT axis as backbone of several top canonical pathways. The central role of AKT regulation towards islet abundance values was further illustrated by its central role in the IPA-generated circular networks.

The late encapsulation effect counterpart (Figure 4b, right panel, Supp. Table 3) encompasses proteins involved in insulin receptor signalling, metabolic pathways and PPARα/RXRa activation, as well as differentiated pancreatic islet markers such as hormones (INS, GCG, PPY, IAPP) and critical components of the hormone processing and secretion (PCSK1, SYT7, IGF2BP3, IGFP5, amongst others). Late encapsulation also regulates epigenetic modifiers with known importance for islet cell fate, such as DNMT1 and DNMT3a. The regulation of DNMT1 was demonstrated previously to alter cell fate maintenance and to promote α-to-β-cell transdifferentiation^36^.

All these results suggest that encapsulation treatment is a driving force promoting β-cell fate by both promoting differentiation at early stages and potentiating the differentiated islet signature during the last, confirming our initial IF results.

### Pathway analysis is suggestive of encapsulation regulating the proteome landscape via integrins

Last, we wanted to characterize the general effects and targets of encapsulation, independent of its action on promoting the pancreatic islet cell fate. For this purpose, we directly compared the encapsulated differentiating cells with their 2D differentiating counterpart (i.e. S5^bead[S0-S5]^ vs S5 as well as S7^bead[S5-S7]^ vs S7, Figure 5a, e).

**Figure 5.**
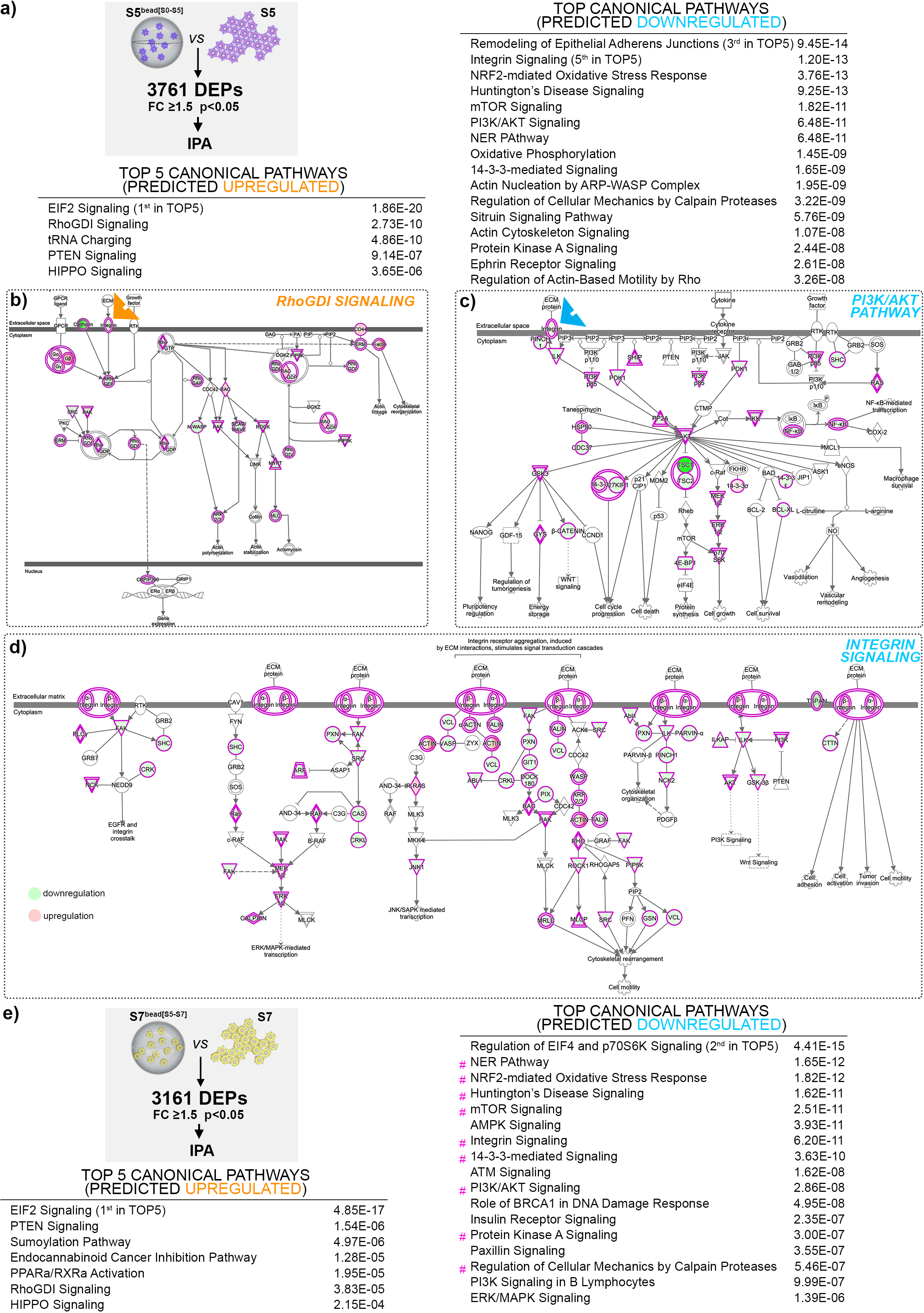
Pathway analysis of the proteome landscape generated by the direct comparison between cells differentiating in alginate capsules and Matrigel differentiated-cells. a) Analysis workflow and IPA-generated tables of the top canonical pathways predicted activated or inhibited at stage 5 of differentiation between S5^bead[S0-S5]^ as compared and S5-cells. b-d) IPA-generated graphical representation of RhoGDI signalling, PI3K/AKT Pathway and Integrin Signalling, e) Analysis workflow and IPA-generated tables of the top canonical pathways predicted activated or inhibited at stage 7 of differentiation between in S7^bead[S5-S7]^ as compared and S7-cells (blue - predicted inhibited, orange – predicted activated, green – observed downregulation, red – observed upregulation, arrow heads point to integrin involvement, magenta # points at shared top pathways between the comparisons). The pathways were generated through the use of IPA (QIAGEN Inc., https://www.qiagenbio-informatics.com/products/ingenuity-pathway-analysis).

The analysis revealed that, besides the already identified pathways involved in islet-cell fate differentiation process, both effects induced the regulation of a wide range of pathways involving integrin signalling (Figure 5, Supp. Figure 5, Supp. Table 4), including RhoGDI Signalling (Figure 5a, b) and PTEN signaling (Figure 5a, Supp. Figure 5a), that were predicted as upregulated, together with predicted downregulation of the PI3K/AKT pathway (Figure 5a, c), the Integrin Signaling itself, (Figure 5a, d), the Regulation of actin-based motility by RHO (Figure 5a, Supp. Fig 5b), and Actin Cytoskeleton Signalling (Figure 5a, Supp. Figure 5c). Interestingly, several of the pro-islet signatures identified above as characterizing the landscapes induced by the encapsulation effects, such as the PI3K/AKT axis or PTEN Signalling, are also shown to be regulated through integrins in our samples (Figure 5c, Supp. Figure 5a). Moreover, we also detected in the top pathways the growth regulating HIPPO pathway (predicted as activated) and the 14-3-3 Signalling pathway (predicted as inactivated) (Supp. Figure 5d, e) characterized by YAP1 downregulation upon encapsulation.

Overall, these results suggested that encapsulation acts during iPSC differentiation through integrins, which probably translate the pressure elicited by the confinement of cells in the alginate matrix into signalling cascades. Of note, these pathways patterns of regulation are consistent with previously published reports^37^, in which their involvement in promoting β-cell differentiation in a different system (micropatterned slides) was proposed, suggesting a general role of the pressure-integrin axis in promoting cell differentiation in confined structures.

## DISCUSSION

Current *in vitro* differentiation protocols for the generation of insulin-producing cells from hPSCs, produce heterogeneous cell populations containing different progenitors and polyhormonal cells^38,39^ that show limited responsiveness to glucose challenges, and are therefore considered immature^4,5^. Nevertheless, transplanting encapsulated hiPSC-derived pancreatic endocrine cells into diabetic mice^40,41,42–45^ concludes the differentiation process and generates functionally mature β-cells, able to maintain glucose homeostasis. The cellular and molecular basis of the *in vivo* process promoting the *in vivo* final β-cell maturation is not known, due to the complex set of systemic interactions acting on the transplanted encapsulated cells.

In this study, our goal was to characterize the specific effects of the encapsulation on the differentiation potential by studying its impact on the differentiating cells proteome fingerprint during either early or late differentiation. In order to eliminate any interference from a possible aggregation/cell clustering effect, we focused on the encapsulation of single cells. This is in contrast with previous studies, which deliberately used clusters to assess the alginate encapsulation effect^29^. We showed here for the first time that encapsulating single cells during pancreas endocrine differentiation promotes a large batch of proteins towards an islet-like proteome landscape, with the cells encapsulated during the later stages of differentiation exhibiting the nearest islet-like profile. Pathway analysis suggests that the effects of encapsulation are transduced via integrins, which translate the cell confinement into intracellular signalling. Of course, the significant islet-profile enhancement of the encapsulated differentiated cells does not imply a significant improvement of their functionality. The acquisition of this parameter might still absolutely require transplantation into living hosts and *in vivo* maturation, hence requiring further investigations.

Our IF results show that cells differentiating in alginate capsules for the entire length of differentiation display an increased percentage of glucagon- and somatostatin-expressing cells, however not insulin. Interestingly, our strategy of defining a narrower window of encapsulation only during the final stages of differentiation (S7^bead[S5-S7]^), *i.e.* from the pancreatic progenitor stage [S5] to maturing islet cell stage [S7]), revealed also a significant increase in the fraction of insulin-expressing cells, clearly suggesting that the timing of encapsulation is decisive for improving hormone-expression. Moreover, our global proteome analysis also revealed that, at the late stages of Matrigel differentiation, the differentiation cocktails promote mostly glucagon expression, while their addition on encapsulated cells will promote all four main islet hormones (insulin, glucagon, somatostatin and PPY). Somatostatin expression is positively modulated regardless of the encapsulation timing, being the hormone with the earliest detected regulation. In contrast, in a different study^29^, late encapsulation of ESC-derived pancreatic progenitor clusters, failed to exhibit an increase in insulin expression, but only glucagon and somatostatin, probably due to the inherently different differentiation protocol employed.

The general impact of the 3D organization for the cell differentiation potential is also supported by the introduction of air liquid interface^1^ or suspension culture^46^ at the last stages of the differentiation protocols. Nevertheless, in these studies^1,4,29^, the changes in the culturing conditions were also combined with clustering of cells aimed at mimicking the *in vivo* developmental niche, hence it was not possible to systematically distinguish between the beneficial effects of the 3D environment and the ones of the intercellular interactions in the clusters.

Regardless of stage, encapsulation did not significantly decrease the fraction of polyhormonal cells; nevertheless, it improved the proportion of insulin-expressing cells co-expressing the key β-cell markers PDX1 and NKX6.1. This is an important result, as previous experiments have revealed that progenitor cells expressing PDX1 and NKX6.1 are the source for functional beta-like cells^47^, while increasing evidence indicates that hPSC-derived polyhormonal cells give rise to glucagon-positive α-like cells^7^.

Our subsequent proteome analyses suggested that encapsulation strongly boosts β-cell fate in a stage-specific manner, by promoting early-stage differentiation and potentiating the islet signature during the later stages. To our knowledge, this is the first comparison by global proteomics of encapsulated and Matrigel differentiating cells. In differentiating encapsulated cells, a large batch of proteins is regulated towards islet cell fate with a significant fraction reaching abundance values similar with islet. We detected a high degree of overlap in the protein populations regulated by encapsulation during early and late regeneration, suggesting that these generally respond to 3D culturing conditions independent of the differentiation stage. We further pinpointed the regulation of growth pathways such as the mTOR signalling, the involvement of the PI3K/AKT axis as well as the INSR signalling in response to encapsulation. The INSR regulatory network is regulated in response to encapsulation during late differentiation specifically, while the PI3K/AKT axis seems to be part of the pathways modulated mostly in cells encapsulated early during differentiation.

It was demonstrated recently that cell confinement is mandatory for endocrine cell specification during development and it negatively regulates the activity of the mechanoresponsive transcription factor YAP1 through a mechanism involving an integrin α5β1-triggered actin cytoskeleton remodelling (F-actin-YAP1-Notch mechanosignaling axis)^37,48^. Similar with these observations, our proteomics data revealed the downregulation of YAP1 in encapsulated cells, coupled with the predicted activation of the Hippo pathway and inactivation of 14-3-3 signalling. Moreover, the pathway analysis on the proteome of cells encapsulated during both early and late differentiation further revealed a large number of top pathways such as RhoGDI Signaling, PTEN Signaling, PI3K/AKT Pathway, Regulation of Actin-Based Motility by RHO, Integrin Pathway, and Actin Cytoskeleton Signalling share an integrin-based regulation. These data suggest that encapsulation acts during hiPSC differentiation acts through integrins by transducing the pressure elicited by confining the cells into alginate matrix to islet-fate promoting signalling cascades. Overall, our results further support the role of confinement and extracellular pressure on positively modulating the differentiation profile of hiPSC. Moreover, we expect that our data will contribute to demultiplexing the intricate outcome of the *in vivo* environment following the transplantation of encapsulated cells, by helping to exclude the protein profiles characterizing the confinement effect inherent to encapsulation.

## MATERIALS AND METHODS

### Cell sources

We used human induced pluripotent stem cells previously generated via episomal reprogramming^31^ from skin fibroblasts collected from a healthy donor. Prior to starting *in vitro* differentiation, the hiPSCs line was enriched for SSEA4^+^ cells using magnetic beads (MACS Miltenyi Biotec), and tested negative for mycoplasma. Human islets were obtained as previously^49^ described from one male and three female deceased donors (age 52-63).

### *In vitro* differentiation

The normal healthy hiPSC line (2 million cells per condition) was differentiated as described previously^31^ following a seven-stage differentiation protocol^1^ in planar culture conditions (on Matrigel-coated plates) up to stage S5 (pancreatic endocrine precursor) and/or stage S7 (maturing beta-cells). 5 million cells were embedded in alginate beads at S0 (hiPSCs) or S5, respectively (Experimental design **Suppl. Fig. 1b**).

### Encapsulation in alginate beads

We used ultra-pure LVG (70 % G and 198 mPas) sodium alginate (batch #BP-0907-02, FMC BioPolymer AS NovaMatrix, Norway) for encapsulation of hiPSC (S0) and S5 cells. Cells were collected using TrypLE Select Enzyme (cat.#12563011, Thermo Fisher), and after viability check and cell counting, were resuspended in 1.8 % alginate in 0.3 M mannitol. The gel beads were formed by using an electrostatic bead machine (Nisco Engineering AG, Switzerland) having a potential difference of 7 kV at a flow of 10 mL/h, and by using a standard nozzle with flat cut tip with an inner diameter of 0.5 mm. Alginate beads were incubated for less than 10 min in gelling solution (50mM CaCl2, 1mM BaCl2 in 0.15M mannitol, 10mM HEPES, pH 7.2)^50^, and rinsed three times in DPBS. The beads were transferred to a 6 cm culture plate and differentiation protocol was continued as described (**Suppl. Fig. 1b**).

### Cell viability and count

Viability and cell count were performed on NucleoCounter NC-200 (ChemoMetec, Denmark) using the Via1-Cassette (cat. no. SKU: 941-0012) with Reagent A100 (cat. no. SKU: 910-0003) and B (cat. no. SKU: 910-0002), as instructed in their protocol for the count of Aggregated Cells A100 and B Assay.

#### Beads processing and IF staining

Alginate beads containing hiPSC-derived cells were fixed in 4% PFA for 1 hour washed in DPBS. For cryosectioning, alginate beads were dehydrated in a sucrose gradient of 10, 20, 30% sucrose. For embedding in Tissue Tek OCT compound (Sakura JP), the beads were placed central in a gelatine capsule (Sanivo Pharma AS) in a plastic mould and frozen. 10 μm sections were obtained by using a cryotome (Leica CM 1950, Leica, DE) and added on gelatine slides. For whole mount, a mean of ten alginate beads (n=6-13) were stained and mounted for each IF staining combination. The following primary antibodies were used: mouse anti-insulin (1/500, I2018, Sigma-Aldrich), guinea-pig anti-porcine insulin (1/400, A056401-2, Dako), mouse anti-porcine glucagon (1/1000, G2654, Sigma-Aldrich), rat anti-somatostatin (1/100, sc-47706, Santa Cruz), guinea-pig anti-PDX1 (1/500, ab47308, Abcam), rabbit anti-PDX1 (1/500, ab47267, Abcam) and rabbit anti-NKX6.1 (1/100, NBP1-82553, Novus). The following secondary antibodies were used: goat anti-guinea-pig A488, goat anti-mouse A546, chicken anti-rat A647, goat anti-mouse A488, goat anti-guinea-pig A647, donkey anti-rabbit A546, and goat anti-mouse A647. The secondary antibodies were all from Molecular Probes (dilution 1/500). DAPI (1/1000, D1306, Molecular Probes) was used to stain the nuclei. The samples mounted in Prolong Diamond Antifade Mountant Media (P36970, Life technologies). Image acquisition was performed using Andor Dragonfly confocal microscope.

### Confocal Imaging

Whole mount beads were imaged using the Andor Dragonfly 5050 (Andor Technologies, Inc) confocal microscope with an 20x dry objective (CFI Plan Apochromat Lambda 20x). Each bead was imaged with 3×3 fields of view, which covered the entire bead. The z-stack were acquired from the top of each bead, and 100 steps of 4 μm with a total of 400 μm depth, which corresponded to the imaging depth. Each image was taken with a high speed iXon 888 Life EMCCD camera with 1024×1024 resolution. For nuclear imaging we used 405 nm laser with intensity of 20-50 % and an exposure time of 100-200 ms. For detecting proteins, laser 488, 546, and 647 were used with laser intensity ranging from 5-20 % and exposure time of 50-200, depending on the antibody.

Furthermore, whole mount and sectioned beads were pictured with a Leica SP5 confocal (Leica) using a 40x immersion objective.

### Image Processing and Analyses

Imaris 9.1.2. (Bitplane AG) was used to analyse the immunofluorescence pictures. A surface mask was used on the DAPI signal, with filters on absolute intensity from 1200 or 1500 to max, quality of more than 100, and size between 150 μm and 10 000 μm with separation of nuclei in clusters of 8 μm in diameter. For the different proteins spot masking was used with quality of more than 100. The MatLab plugin “Find spots close to surface” within 1 μm was used to analyse only spots belonging to a nucleus, which removed the unspecific staining from the analysis. To find colocalizing spots the MatLab plugin “Find colocalizing spots” within 1 μm was used, which gave cells expressing two proteins.

#### Global proteomics analysis

Non-encapsulated hiPSC-derived cells were washed in DPBS and harvested with TrypLE™ Select Enzyme (1X) (12563011, Thermo Fisher Scientific), and collected by centrifugation. Human islets were processed as previously described^31,51^, more specifically here four islet samples representing 200 handpicked equally-sized islets per donor (described above) were combined and mixed to make a homogenous mixture, and 15 μg protein of the mix were divided into three separate samples for downstream TMT 11-plex analysis. Encapsulated cells were lysed directly in the alginate beads, in a buffer containing 8M Urea, 200 mM EPPS pH8.5 and protease inhibitors (Roche complete with EDTA), and sonicated (30 seconds x 3 times at 30% power). Chloroform-Methanol precipitation was performed as previously described^52^. The protein concentration was measured by using a BCA protein assay kit. Samples containing an estimated amount of 15 µg of total protein were processed as described previously^31,51^.

### Data analysis

The mass spectrometry data were analysed as previously described^31,51^. Protein quantitation values were exported for further analysis in Microsoft Excel and GraphPad Prism (version 8). The dataset was uploaded to ProteomeXchange via the PRIDE partner repository with the dataset identifier PXd012704.

The hierarchical clustering was performed with GeneSpring 14.9.1 GX software (Agilent), with clustering on both entities and conditions by using Squared Euclidian distance metric and Ward’s linkage rule. The pathway analyses were generated by QIAGEN’s Ingenuity Pathway Analysis program (IPA®, QIAGEN Redwood City, www.qiagen.com/ingenuity)^53^ as previously described^31,51^, here using 35 molecules/network; 25 networks/analysis for generating the interaction networks.

### Statistical analysis

Statistical analysis on the proteomics data was tested using unpaired two-tailed Student’s *t*-test, and a *p*-value of ≤0.05 was considered significant. Two-way ANOVA with Tukey's multiple comparisons test were used to compare between the groups for number of cells positive for the different markers by IF staining. This analysis was performed using GraphPad Prism v8.1.2.

### Data availability

The materials, methods and data sets that support the findings of this study are available upon request from the corresponding author (S.C.).

### Ethical Statement

The reported experimental protocols were approved by the Regional Committee of Medical and Health Research Ethics, for hiPSCs (REK 2010/2295) and for human islets (REK 2011/426), and all methods were carried out in accordance with the Helsinki Declaration. Informed consent was obtained from relatives for organ donation and for its use in research (human islets).

## Acknowledgements

We thank A.H. Knudsen for technical help and H.A. Dale for Imaris assistance. The confocal imaging was performed at the Molecular Imaging Center (MIC), Department of Biomedicine, University of Bergen. Human islets were provided through the Islet Distribution Program at University of Oslo to H.S. The authors are grateful to the human islet isolation team in Oslo, Norway. We would also like to thank the Gygi Lab and Taplin Facility at Harvard Medical School, particularly Dr. Steven P. Gygi for the use of his mass spectrometers. This work was funded with grants from the Research Council of Norway (NFR 247577 and 251041) and the Novo Nordisk Foundation (NNF15OC0015054) to S.C. J.A.P. is funded by NIH/NIDDK grant DK098285. H.R. is supported by grants from Bergen Forskningsstiftelse (BFS2014REK02) and the Western Norway Regional Health Authority (Bergen Stem Cell Consortium).

## Author contributions

H.V. performed the differentiation, sample preparation for proteomic analyses, immunofluorescence staining, analysed data and wrote the manuscript; H.V. and S.C. analysed the proteomics data; T.A.L performed immunofluorescence imaging and analyses, edited and revised the manuscript; B.L.S and L.G. established the encapsulation protocol; S.A. and H.S. generated human islet preparations; J.A.P. performed the TMT-labelling experiment and mass spectrometry analysis; H.R. provided the iPSC cell line and access to the iPS facility; H.S. and H.R. edited the manuscript; L.G. and S.C. conceived the experiments, interpreted the observations, wrote and revised the manuscript. All authors approved the final version of the manuscript.

