## Supplementary material for "Encapsulation boosts islet-cell signature in differentiating human induced pluripotent stem cells via integrin signalling": Supp. Figure

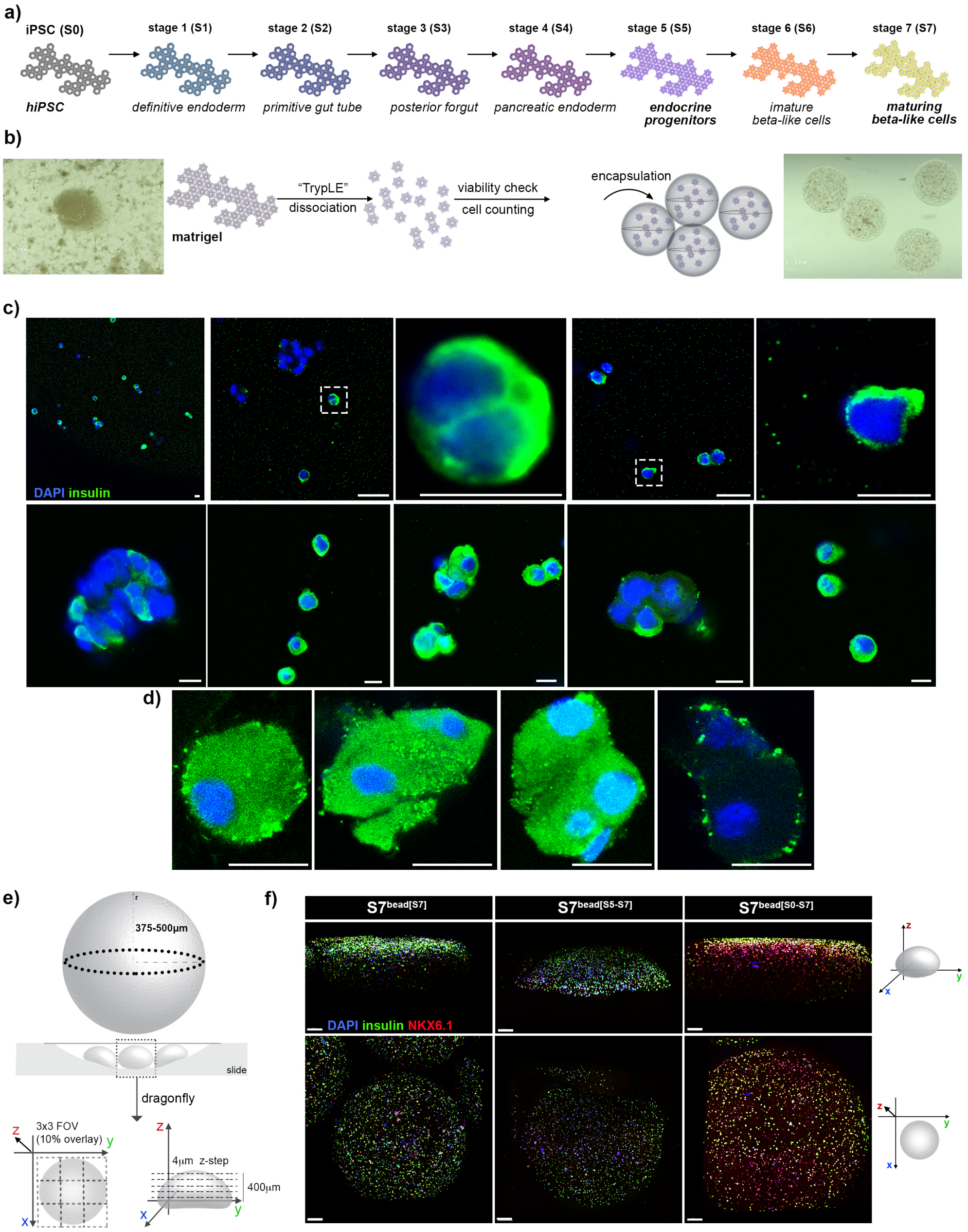


**Supplemental Figure 1.** a) Scheme depicting the seven stages of the hiPSC differentiation protocol. b) Experimental design of the encapsulation procedure. c) High magnification confocal images of encapsulated cells inside alginate capsules following whole mount immunofluorescence (insulin – green, DAPI – blue). d) High magnification confocal images of encapsulated cells following alginate capsule cryosectioning and immunofluorescence staining for insulin (green) and DAPI (blue). e) Scheme depicting the procedure for the imaging of the encapsulated cells with the Dragonfly confocal, f) 3D reconstructions of insulin (green), NKX6.1 (red), DAPI (blue) immunofluorescence on whole alginate beads containing the three distinct populations analyzed. Scale bar: c, d - 10µm, f - 150µm.

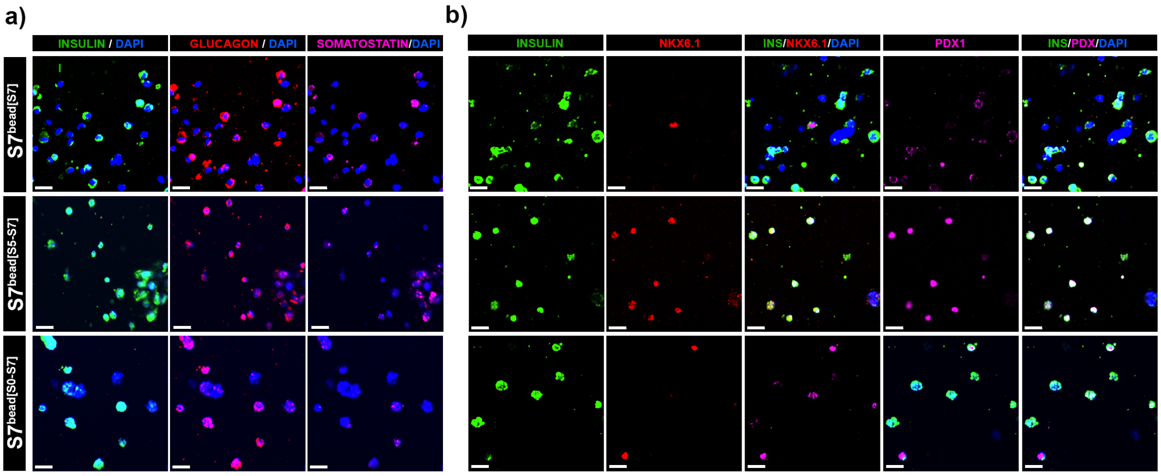


**Supplemental Figure 2.** a) Immunofluorescence staining of insulin (green), glucagon (red) and somatostatin (purple), DAPI (blue) of the three distinct populations analyzed. b) Immunofluorescence staining of insulin (green), NKX6.1 (red) and PDX1 (purple), DAPI (blue) of the three distinct populations analyzed. Scale bars: 20µm, gamma correction 0.4.

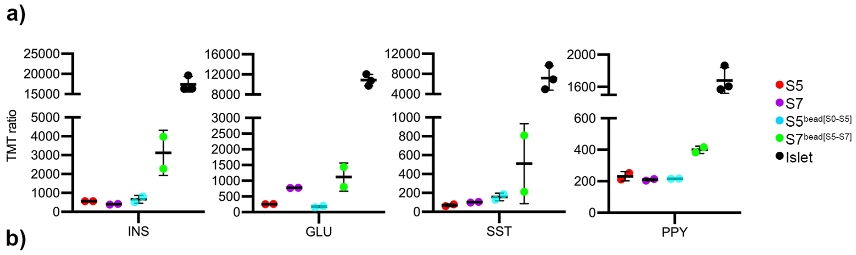


b)

| Comparison | Discriminated by |
|---|---|
| S5-cells/I vs S7-cells/I | Stage/Differentiation cocktail |
| S5-cells/I vs S5bead[S0-S5] | Encapsulation |
| S5-cells/I vs S7bead[S5-S7] | Stage/Differentiaton cocktail & Encapsulation |
| S7-cells/I vs S7bead[S5-S7] | Encapsulation |

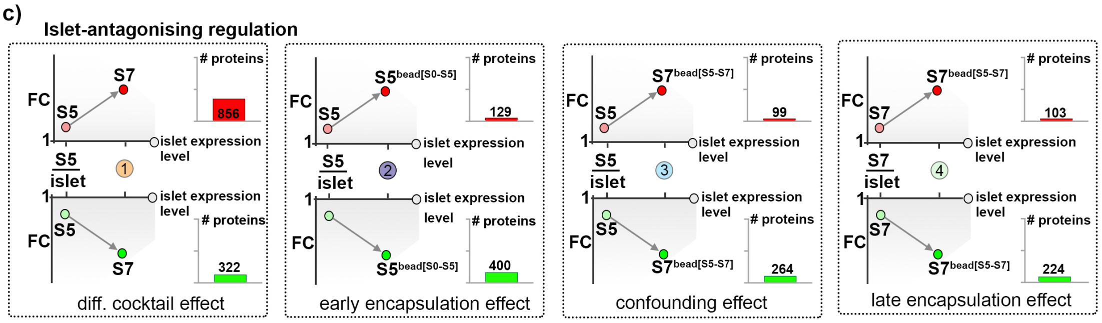


**Supplemental Figure 3. a)** The TMT-ratios of the four main pancreatic hormones in the five conditions analyzed. Graphs data are shown as mean ± SEM. b) Table listing the biological process responsible for the difference between the samples compared. Arrows depict the generic prerequisite direction of regulation for group inclusion. c) The number of proteins showing a dynamic of regulations compatible with an islet-antagonizing pattern in response to each of the four effects considered.

a) **DIFFERENTIATION COCKTAIL EFFECT**

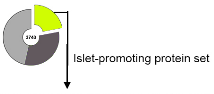


TOP PREDICTED UPSTREAM **TRANSCRIPTION** REGULATORS

| NAME | activation z-score | prediction |
|---|---|---|
| MYC | -3.904 | Inhibited |
| SRF | -3.197 | Inhibited |
| REL | -3.023 | Inhibited |
| SREBF1 | -2.923 | Inhibited |
| SREBF2 | -2.734 | Inhibited |
| RB1 | 3.368 | Activated |
| PML | 2.186 | Activated |
| RBL1 | 2.000 | Activated |
| NEUROG1 | 2.000 | Activated |

b) **DIFFERENTIATION COCKTAIL EFFECT**

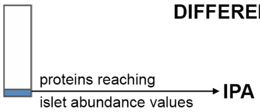


| TOP CANONICAL PATHWAYS | |
|---|---|
| Tumoricidal Function of Hepatic NK Cells | 1.99E-04 |
| Mitochondrial Dysfunction | 4.74E-04 |
| IGF-1 Signaling | 5.14E-04 |
| Tight Junction Signaling | 1.18E-03 |
| Valine Degradation I | 1.32E-03 |
| Isoleucine Degradation | 1.01E-04 |

TOP PREDICTED UPSTREAM REGs

| NAME | p-value |
|---|---|
| HNF4A | 4.84E-07 |
| TP53 | 3.26E-06 |
| KRAS | 9.54E-06 |
| MMP3 | 6.41E-05 |
| E2F1 | 2.17E-04 |

c) **CONFOUNDING EFFECT**

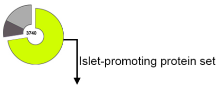


TOP PREDICTED UPSTREAM **TRANSCRIPTION** REGULATORS

| NAME | activation z-score | prediction |
|---|---|---|
| DNAJB6 | -2.412 | Inhibited |
| FOXO4 | -2.138 | Inhibited |
| HOXC8 | -2.000 | Inhibited |
| RB1 | 3.704 | Activated |
| CDKN2A | 2.928 | Activated |
| NFE2L2 | 2.885 | Activated |
| HES3 | 2.646 | Activated |
| KLF15 | 2.641 | Activated |
| SOX4 | 2.621 | Activated |
| NEUROD1 | 2.590 | Activated |
| TFEB | 2.575 | Activated |
| MAF | 2.408 | Activated |

d) **CONFOUNDING EFFECT**

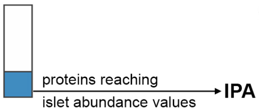


| TOP CANONICAL PATHWAYS | |
|---|---|
| Protein Ubiquitination Pathway | 4.48E-15 |
| EIF2 Signaling | 4.74E-09 |
| NER Pathway | 2.59E-07 |
| Estrogen Receptor Signaling | 1.05E-06 |
| Hereditary Breast Cancer Signaling | 2.32E-06 |
| PTEN signaling | 4.23E-06 |
| mTOR signaling | 4.83E-06 |
| PI3K/AKT Signaling | 8.26E-06 |
| Molecular Mechanism of Cancer | 8.68E-06 |
| Reg. of EIF4 and p70S6K Signaling | 9.06E-06 |
| Insulin Receptor Signaling | 1.69E-05 |

TOP PREDICTED UPSTREAM REGs

| NAME | p-value |
|---|---|
| HNF4A | 7.61E-18 |
| CST5 | 3.11E-10 |
| MYC | 1.49E-09 |
| RICTOR | 3.42E-09 |
| MYCN | 6.68E-09 |

e) **EARLY ENCAPSULATION EFFECT**

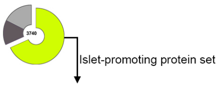


TOP 5 PREDICTED UPSTREAM **TRANSCRIPTION** REGULATORS

| NAME | activation z-score | prediction |
|---|---|---|
| TCF7L2 | -4.814 | Inhibited |
| E2F1 | -4.529 | Inhibited |
| MYC | -4.224 | Inhibited |
| CTNNB1 | -3.928 | Inhibited |
| FOXM1 | -3.926 | Inhibited |
| NUPR1 | 3.796 | Activated |
| CDKN2A | 2.635 | Activated |
| PML | 2.621 | Activated |
| PDX1 | 2.512 | Activated |
| MYCN | 2.369 | Activated |

f) **EARLY ENCAPSULATION EFFECT**

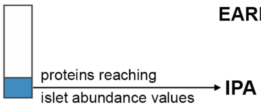


| TOP CANONICAL PATHWAYS | |
|---|---|
| Regulation of eIF and p70S6K Signaling | 8.05E-09 |
| EIF2 Signaling | 3.29E-07 |
| FAK Signaling | 3.33E-07 |
| Cell Cycle Reg. by BTG Family Proteins | 5.71E-07 |
| tRNA Charging | 9.75E-07 |
| mTOR Signaling | 2.45E-06 |
| Rac Signaling | 2.77E-06 |

TOP PREDICTED UPSTREAM REGs

| NAME | p-value |
|---|---|
| HNF4A | 1.91E-15 |
| RICTOR | 5.39E-09 |
| MMP3 | 7.22E-08 |
| MYC | 3.22E-06 |
| CST5 | 5.73E-06 |

g) **LATE ENCAPSULATION EFFECT**

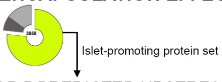


TOP 5 PREDICTED UPSTREAM **TRANSCRIPTION** REGULATORS

| NAME | activation z-score | prediction |
|---|---|---|
| E2F1 | -4.671 | Inhibited |
| MYC | -4.067 | Inhibited |
| FOXM1 | -3.807 | Inhibited |
| MITF | -3.803 | Inhibited |
| TBX2 | -3.273 | Inhibited |
| NUPR1 | 4.279 | Activated |
| CDKN2A | 3.899 | Activated |
| RBL1 | 2.896 | Activated |
| RB1 | 2.669 | Activated |
| NCOA1 | 2.632 | Activated |

h) **LATE ENCAPSULATION EFFECT**

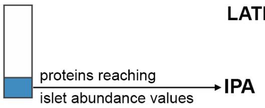


| TOP CANONICAL PATHWAYS | |
|---|---|
| Protein Ubiquitination Pathway | 1.14E-15 |
| EIF2 Signaling | 3.50E-13 |
| NER Signaling | 3.48E-09 |
| Regulation of eIF and p70S6K Signaling | 5.01E-06 |
| mTOR Signaling | 9.50E-06 |

TOP PREDICTED UPSTREAM REGs

| NAME | p-value |
|---|---|
| HNF4A | 7.61E-18 |
| CST5 | 3.11E-10 |
| MYC | 1.49E-09 |
| RICTOR | 3.42E-09 |
| MYCN | 6.68E-09 |

**Supplemental Figure 4.** Tables depicting the top predicted upstream transcription regulators in response to Differentiation Cocktail Effect. b) Pathways analysis of the proteins reaching abundance levels indistinguishable from the ones detected in native human islets in response to Differentiation Cocktail Effect, c) Table depicting the top predicted upstream transcription regulators in response to Confounding Effect, d) Pathways analysis of the proteins reaching abundance levels indistinguishable from the ones detected in native human islets in response to Confounding Effect, e) Table depicting the top predicted upstream transcription regulators in response to Early Encapsulation Effect, f) Pathways analysis of the proteins reaching abundance levels indistinguishable from the ones detected in native human islets in response to Early Encapsulation Effect, g) Table depicting the top predicted upstream transcription regulators in response to Late Encapsulation Effect, h) Pathways analysis of the proteins reaching abundance levels indistinguishable from the ones detected in native human islets in response to Late Encapsulation Effect.

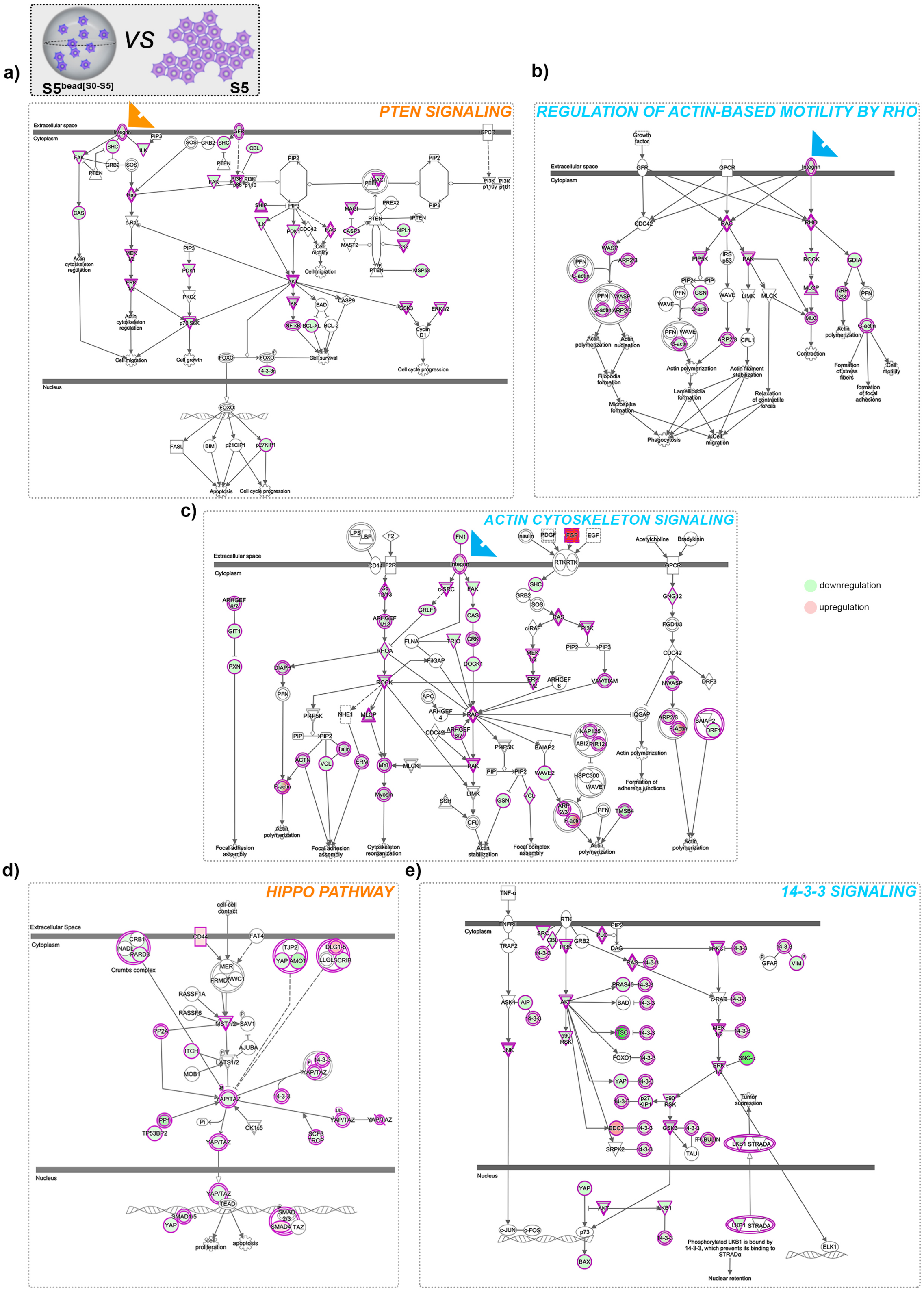


**Supplemental Figure 5.** Graphical representations of selected top activated canonical pathways and their observed protein occupancy characterizing the direct comparison between S5-cells differentiated on Matrigel (S5) and S5-cells differentiated in alginate capsules ($S5^{bead[S0\text{-}S5]}$). a) PTEN Signaling, b) Regulation of Actin-Based Motility By Rho Signaling, c) Actin Cytoskeleton Signaling, d) Hippo Signaling, e) 14-3-3 Signaling. (blue - predicted inhibited, orange – predicted activated, green – observed downregulation, red – observed upregulation, arrow heads point to integrin involvement). The networks were generated through the use of IPA (QIAGEN Inc., https://www.qiagenbio-informatics.com/products/ingenuity-pathway-analysis).
